# Functions of farmers’ preferred tree species and their potential carbon stocks in southern Burkina Faso: implications for biocarbon initiatives

**DOI:** 10.1101/344408

**Authors:** Kangbéni Dimobe, Jérôme E. Tondoh, John C. Weber, Jules Bayala, Karen Greenough, Antoine Kalinganire

**Author notes:** **Corresponding author** (KD).

## Abstract

The success of terrestrial carbon sequestration projects for rural development in sub-Saharan Africa lies in the (i) involvement of local populations in the selection of woody species, which represent the biological assets they use to meet their daily needs, and (ii) information about the potential of these species to store carbon. Although the latter is a key prerequisite, there is very little information available. To help fill this gap, the present study was undertaken in four pilot villages (Kou, Dao, Vrassan and Cassou) in Ziro Province, south-central Burkina Faso. The objective was to determine carbon storage potential for top-priority woody species preferred by local smallholders. We used (i) participatory rural appraisal consisting of group discussions and key informant interviews to identify priority species and functions, and (ii) landscape assessment of carbon stocks in the preferred woody species. Results revealed over 79 priority tree and shrub species grouped into six functions, of which medicine, food and income emerge as the most important ones for the communities. For these functions, smallholders overwhelmingly listed *Vitellaria paradoxa*, *Parkia biglobosa*, *Afzelia africana*, *Adansonia digitata*, *Detarium microcarpum*, and *Lannea microcarpa* among the most important tree species. Among the preferred woody species in Cassou and Kou, the highest quantity of carbon was stored by *V. paradoxa* (1,460.6 ±271.0 kg C ha^−1^ to 2,798.1±521.0 kg C ha^−1^) and the lowest by *Grewia bicolor* (1.6±1.3 kg C ha^−1^). The potential carbon stored by the preferred tree communities was estimated at 5,766.2 Mg C ha^−1^ (95% CI: 5,258.2; 6,274.2 Mg C ha^−1^) in Kou and 6,664.0 Mg C ha^−1^ (95% CI: 5,810.2; 7,517.8 Mg C ha^−1^) in Cassou. The findings of this study will help design data-based development of biocarbon projects, which are rare in the West African Sahel despite being considered as one of the most impactful climate change resilient strategies.

## Introduction

In the West African Sahelian and Sudanian agro-ecological zones, parkland agroforests are socio-ecological systems that integrate trees, crops and livestock. They play key roles in the functioning of agro-ecological landscapes, delivering essential goods and many ecosystem services that sustain smallholder farmers and pastoralist livelihoods [1-8]. Delivery of provisioning ecosystem services, including food, fodder and fuel wood, contributes greatly to rural communities’ daily needs. Income from these products helps to improve livelihoods and build resilient socio-ecological systems in the face of ongoing climate change and variability [9]. These integrated tree-crop-livestock systems are subject to severe degradation through deforestation due to both climate change [10] and unsustainable land management practices, such as overgrazing and wood cutting [11-12]. This poses immediate threats to smallholders’ sustainability and coping abilities in confronting the adverse impacts of climate change. Land degradation leads to a reduction of vegetation cover, species richness and abundance. The corollary is an increase in soil erosion, depleting soil nutrients, including soil organic carbon. Thus, the low standing biomass of degraded land is associated with low soil carbon and diminished productive properties, weakening the resilience of farming systems and that of people making their living from these systems [13].

Restoring agro-ecological functions for increased productivity and resilience requires climate smart land uses [14-16]. By encompassing a set of land cover options and management practices that increase greenhouse gas (CO_2_) absorption and biocarbon stocks, these land uses can also help mitigate climate change by reducing overall concentration of CO_2_ in the atmosphere [13]. Biocarbon projects are among the options which have been promoted in the framework of the Kyoto Protocol and more recently through REDD+ initiatives. The latter intend to contribute to local development by generating carbon-based incomes for smallholders through carbon markets. Co-benefits include supporting ecosystem services like improved soil fertility, and provisioning services like food and income [13, 17-18]. Biocarbon projects are diverse in scope, and a large share of World Bank Biocarbon Fund investments go into environmental restoration of degraded lands (50.5%), fuelwood production (23%) and timber production (20%) [16].

The effectiveness of biocarbon initiatives to improve smallholders’ livelihoods through carbon finance has been challenged, particularly in the Sahel [ 19], because of many bottlenecks, including (i) the dearth of empirical data on the potential of farmers’ preferred tree species to store carbon, (ii) the long time it takes for biocarbon projects to become economically viable and profitable, (iii) the complexity of access to carbon markets, (iv) the uncertainty about future climate, and (v) low carbon prices in international markets. Farmer-managed natural regeneration (FMNR), a land management practice many Sahelian farmers use to rehabilitate their degraded lands, could be the foundation for biocarbon initiatives [20-21]. However, the data and knowledge gap (carbon storage potential) must be bridged in a participatory way before best-fit options for Sahelian biocarbon initiatives may be scaled-up.

The objectives of the present study were to determine priority species, functions, and carbon storage potential of woody species that local communities deemed top-priority or very useful. This work was carried out within the framework of the Building Biocarbon and Rural Development in West Africa (BIODEV) project implemented in Burkina Faso, Guinea-Conakry, Mali and Sierra Leone. The overall goal of the project was to demonstrate the multiple developmental and environmental benefits that result from a high value biocarbon approach to climate change and variability in large landscapes [22]. The current study falls under “Agroforestry and farm interventions” that aimed at increasing the adoption of agroforestry and other carbon-enriching farm practices that meet beneficiaries’ priority needs and address climate change issues.

## Materials and methods

### Study area

The study sites were in the following villages: Kou, Dao, Vrassan and Cassou (Fig 1), located in Ziro Province (11°16’N to 11 °45’N and 2°10’W to 1°48’W) in south-central Burkina Faso. These villages were selected because local populations have close interactions with the Cassou Forest [12, 23].

**Fig 1.**
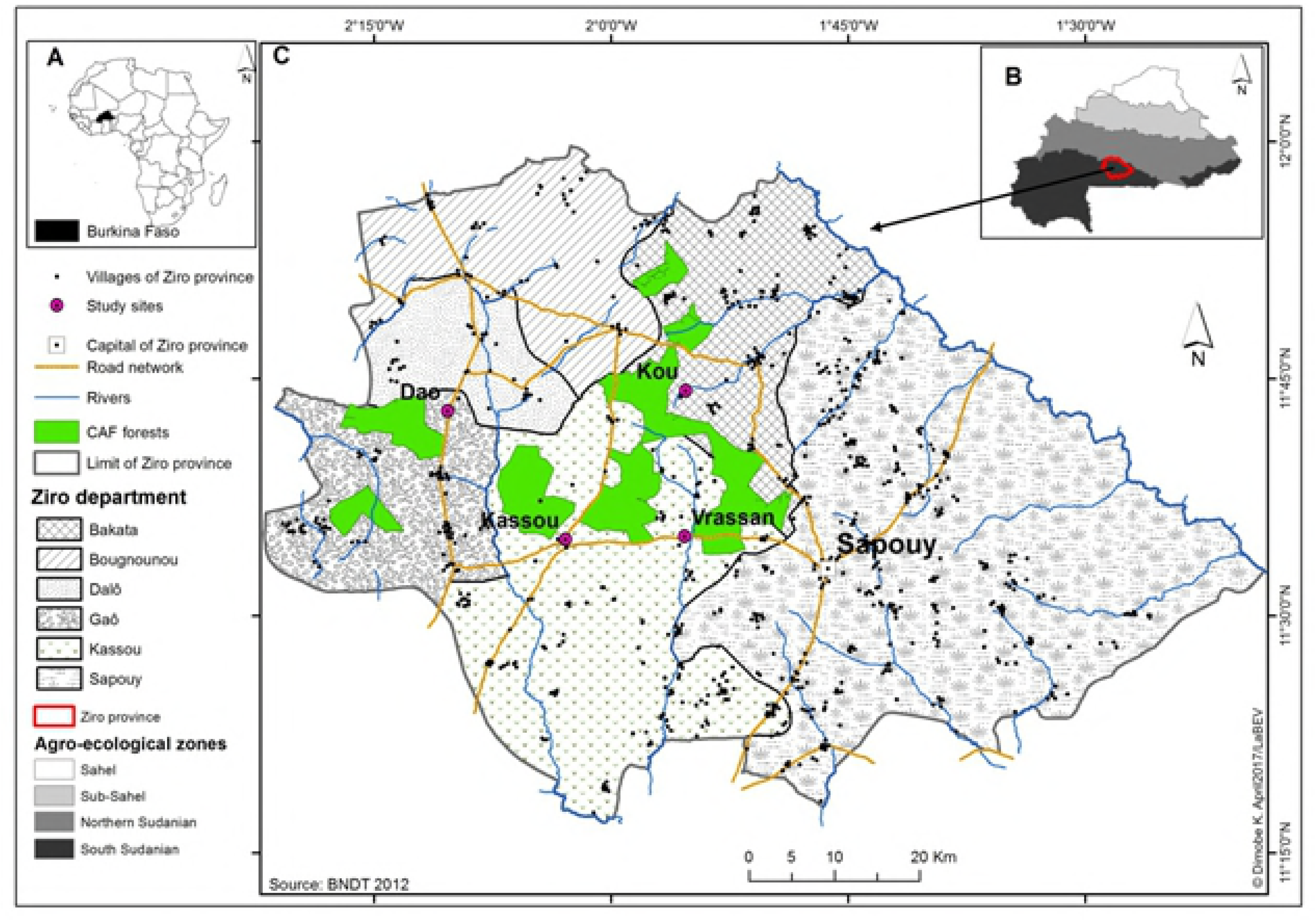
Location of the study area. (A) the location of Burkina Faso in Africa, (B) the position of province of Ziro in Burkina Faso, and (C) the province of Ziro with study sites.

The Ziro province covers 5,291 km^2^ and is characterized by a low-relief topography with a mean altitude of 300 masl. Phytogeographically, this province is located within the south-Sudanian agro-ecological zone [24], which receives higher rainfall in the country, 800-1,000 mm annually. The rainfall follows a unimodal pattern that lasts for six months (May through October), and temperatures in the province range from 30°C on average to peaks of 40°C during the dry season. Twenty-five villages surround the 30,000 ha of Cassou forest, comprising a mixture of mostly dry natural forest and tree savannah vegetation types. Main soil types are silt-clay cambisols, sandy lixisols, and loamy ferric luvisols [25].

The villages of Dao and Kou belong to the departments of Gao and Bakata, respectively, while Cassou and Vrassan fall in Cassou department. The province has one of the highest average rural population densities in the country, with an estimated 30 inhabitants per km^2^ [26-27]. The dominant production systems are cereals rotated with tubers, and animal husbandry.

Supervised by a private organization, *Chantiers d’Aménagements Forestiers* (CAF, Community Forest Management), local populations jointly implement a management plan for the protected Cassou forest. The CAFs were created by the government in this region to ensure a sustainable supply of commercial fuel wood, construction poles, and other forest products to the nearby cities of Ouagadougou and Koudougou [23], and thereby contribute to local community livelihoods. The CAFs’ primary challenge lies in the sustainable management of the forest.

## Data collection

### Priority woody species

The study used a participatory approach within smallholder communities to assess priority tree functions and species and collect information about products and services provided by them [28]. The process, carried out in two stages in the four study sites (Fig 1), consisted of gendered group discussions held in each village from July through August 2013, one for adult men and one for women, followed by individual interviews with key informants in each village. A pilot survey conducted in the four sites (Cassou, Dao, Kou and Vrassan) with 30 respondents randomly selected per site determined the number of respondents to be interviewed. The sample size was determined following the normal approximation of the binomial distribution of Dagnelie [29]:
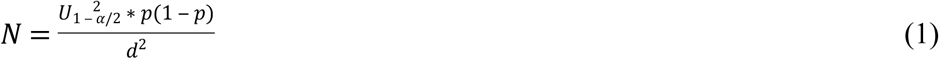

where *N* = the estimated sample size; *U* = the value of the normal random variable (1.96 for α = 0.05), *p* = the proportion of respondents who knew and used the preferred woody species; and *d =* the error margin assumed to be 5%.

The pilot survey revealed that 91% of respondents in Cassou and Kou, 90% in Dao and 89% in Vrassan had a good knowledge and used the preferred woody species. Thus, for an error margin of 8%, 49, 50, 55 and 59 people were randomly selected for the participatory group discussions in Cassou, Kou, Dao and Vrassan, respectively (Table 1).

**Table 1.**
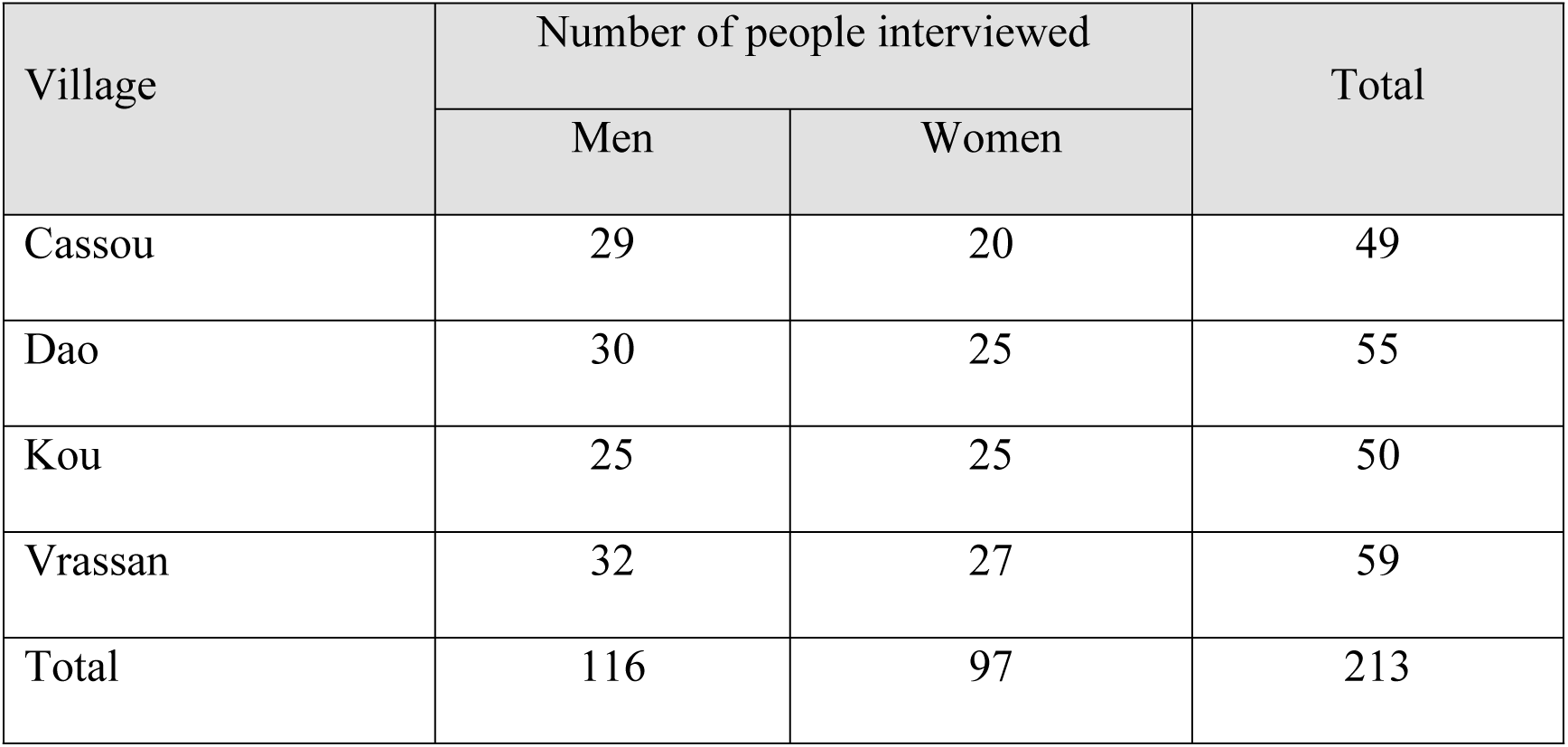
Number of localities in which surveys and questionnaires were administered to select priority woody species in the study area (Cassou, Dao, Kou and Vrassan) in Ziro province of Burkina Faso

Discussions, surveys, and interviews were undertaken with the help of field assistants who understood and spoke the local language.

During the group discussions, information concerning products and services provided by woody species was obtained in four steps. First, the groups of men and women generated two lists of woody species they considered important, along with the products and services provided by each species. Secondly, each group discussed and reached a consensus on six priority tree functions, namely food, sale, medicine, fodder, energy, and craft. Thirdly, to qualify the functions of each species the groups assigned scores ranging from 0 to 3, with 0 corresponding to “not useful” and 3 to “very useful”. The importance of each species was estimated by calculating the sum of the six tree function scores, referred to hereafter as species importance value. The species importance values were then used to identify the priority species of the different social groups in each village. The importance of each function was determined by calculating the sum of its scores across all species. Lastly, the two gender groups came together to thoroughly review the priority species that each identified and reach a consensual list of priority woody species for the village. Thus, the list of priority species appears to reflect the local communities’ assessment and the species’ contributions to improving their livelihoods.

The vernacular names of woody species supplied by respondents in this study were crosschecked with previous studies: e.g. [30-31], as recommended by Nolan and Robbins [32]. The scientific names of cited woody species and their authorities were validated using the website of West African Plants (http://www.westafricanplants.senckenberg.de/root/index.php) and Vascular Plants of Burkina Faso [30].

### Above-ground biomass and carbon stocks determination

Carbon stocks of preferred woody species were calculated using above-ground biomass (AGB) field data collected in the two contrasting landscapes of Cassou and Kou villages spanning agroecological landscapes and portions of Cassou Forest. Each sampling site covered 100 km^2^ (10 km x 10 km) where the Land Degradation and Surveillance Framework (LDSF) was applied [33]. A hierarchical stratified random sampling design was used in the sampling site consisting in 16 clusters of 100 ha each. Each cluster was composed of 10 plots of 0.1 ha established randomly (Fig 2) and further divided into four subplots of 0.01 ha of 5.64 m radius as per a standard layout (Fig 2b).

**Fig 2.**
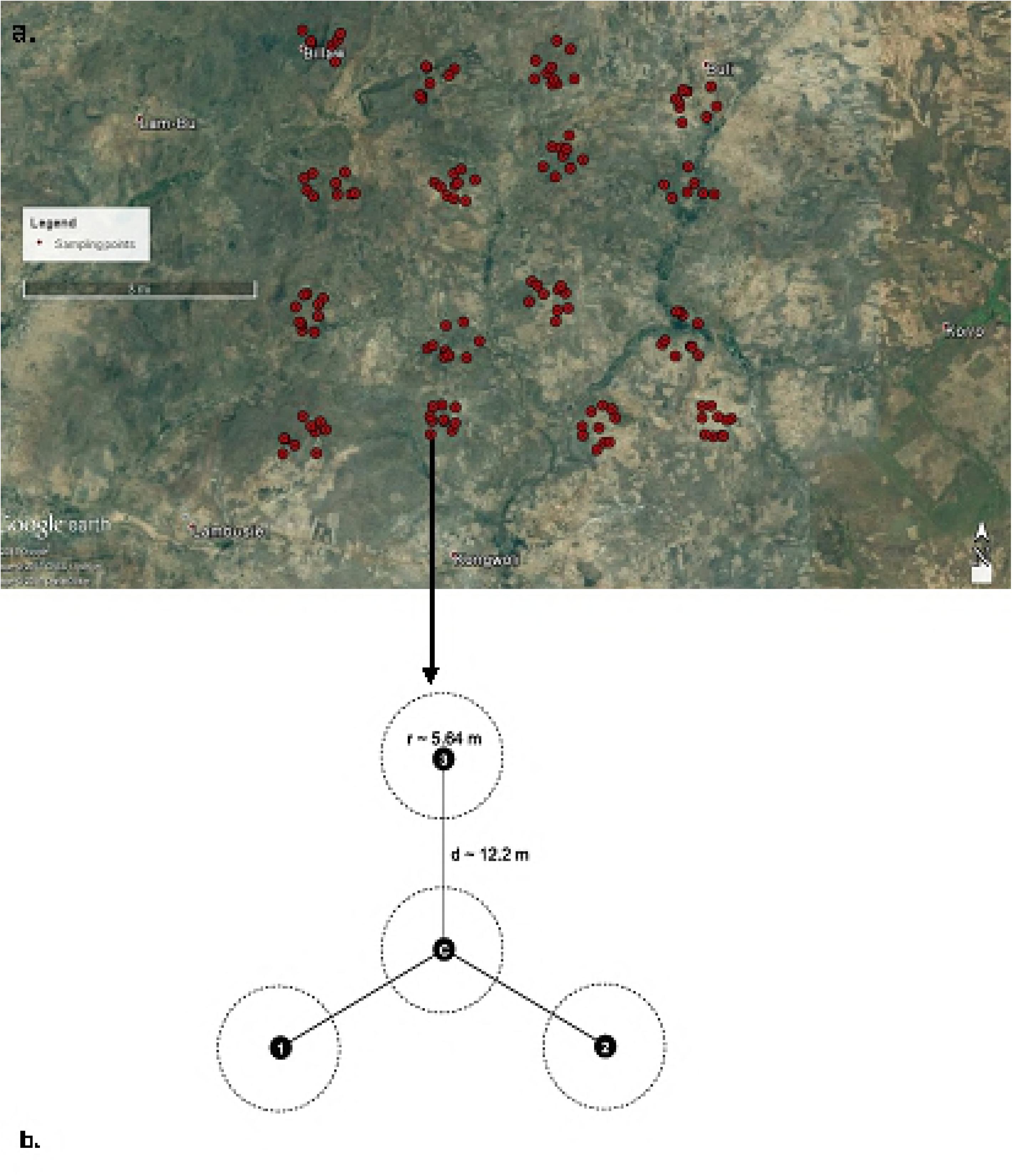
LDSF hierarchical sampling. (a) plots grouped in 16 clusters within the landscape (10 km x 10 km), and (b) sampling layout of subplots c, 1, 2, 3.

The central position of subplot c was located by the Global Positioning System (GPS), and 12.2 m measured to the upper slope position to mark the center of the subplot 3. Subplots 1 and 2 were located by offsetting 120 and 240 degrees from subplot 3. In each subplot, individual trees were sampled for measurement using the T-square method, which is recommended for plant communities where individuals are randomly distributed [34-36]. The diameter at breast height (dbh) over bark for each tree ≥ 5 cm was measured with a diameter tape at 1.3 m above ground level. For trees forking below 1.3 m, the diameter of all ramifications was measured and the quadratic mean diameter (root-mean-squared) was calculated as:
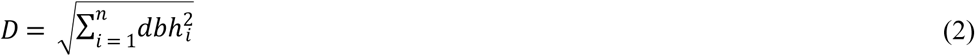

where *D* = the quadratic mean diameter; *dbhi* = the diameters of the measured stems.

Tree heights were estimated using a pole or a clinometer. To estimate aboveground tree biomass (AGB), tree species with height ≥ 3 m were considered as containing the greater portion of aboveground biomass [37-38]. Since allometric equations are not available for most of the species, AGB was estimated using the generalized allometric model equation 4 of Chave et al. [39]:
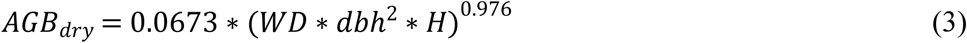

Where *H* = height (m); dbh = diameter at breast height; *WD* = wood density (g cm^−3^).

The *getWoodDensity* function from the BIOMASS package was used to assign a wood density value to each taxon using the global wood density database as a reference [40-41]. By default, *getWoodDensity* assigns to each taxon a species- or genus-level average if at least one wood density value in the same genus as the focal taxon is available in the reference database [42]. To upscale from the tree level to the site level (100 km^2^ or 10,000 ha), the predicted AGB was first calculated at the cluster scale (100 ha) and averaged based on the total number of plots. The AGB (Mg ha^−1^) was further upscaled to the study site level (10,000 ha) by applying the surface expansion factor (100). For carbon stock determination, the AGB (Mg ha− ^1^) was converted by applying a carbon conversion factor of 0.5 [43-44].

### Data processing and statistical analyses

Statistical analyses were done in the R statistical software package, version 3.3.2 [45]. Statistical assumptions were explored visually as proposed by Zuur and colleagues [46]. We first checked for normality among the different variables using the Shapiro–Wilk normality test. To identify farmers’ preferences for trees and shrubs based on their functions in households, we next processed the data collected from the participatory group discussions to derive key variables, including the number of very useful or top-priority species, total species richness per gendered group, and tree function scores. We also tested the influence of the villages on the number of very useful species, species richness and tree function scores using village as a random factor.

As some data were not normally distributed, Wilcoxon Rank Sum test was used for probing the statistical differences between two variables and Kruskal-Wallis Rank Sum test was used for more than two variables.

Two Principal Component Analyses were performed using the FactoMineR package to (i) determine the relationship between tree functions (Medicine, Food, Sale, Energy, Fodder) and the priority woody species identified by local communities, and (ii) assess the relationships between tree functions (Medicine, Food, Sale, Energy, Fodder) and the measured variables (scores of gender groups for each function) across the four villages.

The “*computeAGB*” function from the BIOMASS package has been used to calculate the AGB. Statistical analyses were performed at a significance level of 0.05.

## Results

### Floristic diversity and preferences for tree functions

In total, 79 woody species representing 65 genera and 26 families were listed by respondents in the four villages as important tree and shrub species (See S1 Table). The most represented families were Fabaceae-Mimosoideae (9 species), Fabaceae-Caesalpinioideae (9 species), followed respectively by Combretaceae (8 species), Anacardiaceae, Rubiaceae (6 species), Moraceae and Malvaceae (5 species). Among these important species, 18 were identified as top-priority species (Fig 3). *Afzelia africana* (100%: in all the 4 villages), *Bombax costatum* (100%), *Parkia biglobosa* (100%), *Pterocarpus erinaceus* (100%), *Tamarindus indica* (100%), *Vitellaria paradoxa* (100%) were mentioned by all respondents.

**Fig 3.**
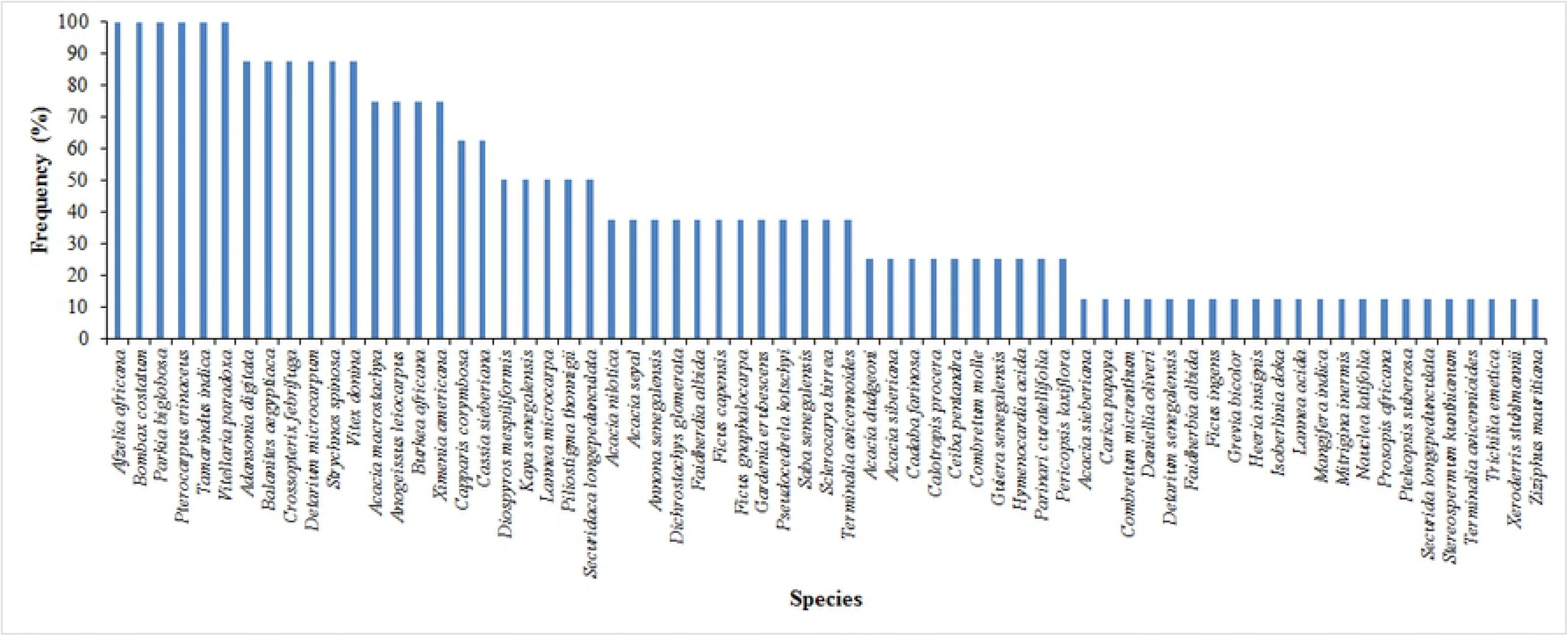
Frequency curve of species across the study villages (Cassou, Dao, Kou and Vrassan) in Ziro province of Burkina Faso.

Respondents listed “Medicine” (96%), “Food” (94%), “Sale” (87%), “Energy” (56%), “Fodder” (45%) and “Craft” (4%) as the main priority functions for tree and shrub species in the study area. The villages did not influence significantly the number of very useful species (*p* = 0.58), species richness (*p* = 0.36) and tree function scores (*p* = 0.83) provided by the interviewees (Fig 4a). This means that the respondents in the four villages valued the species the same way giving function scores of similar magnitudes to the very useful species.

**Fig 4.**
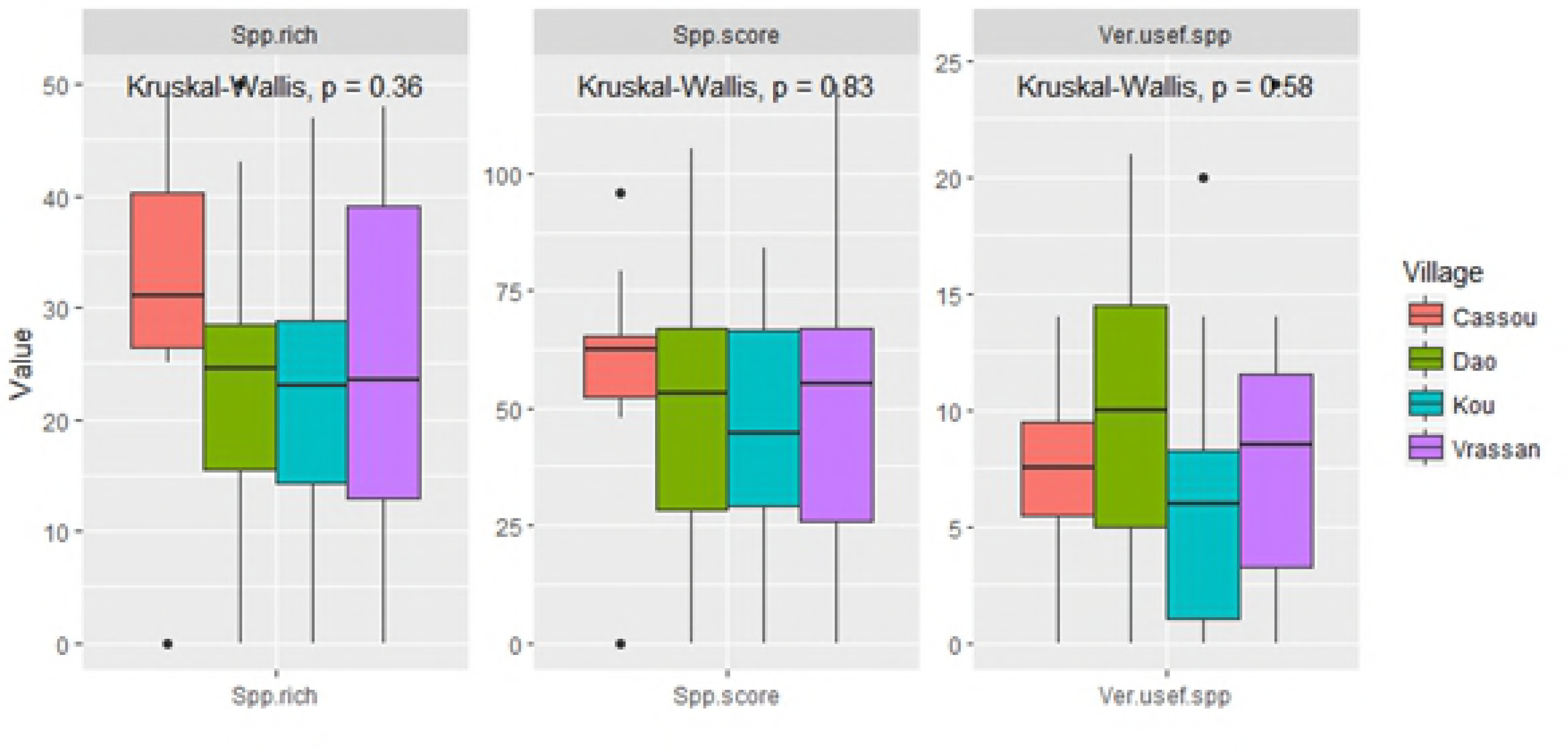
Abundance of very useful tree species, species richness and total score of tree function in Ziro province of Burkina Faso per (a) pilot village (Cassou, Dao, Kou and Vrassan) and (b) per tree function. Spp.rich= species richness; Spp.score= total score of tree function; Ver.usef.spp= very useful tree species

In contrast, the number of top priority woody species (*p* <0.001), total species richness (*p* < 0.00001) and total score (*p* < 0.001) differed significantly among the functions (Fig 4b). “Medicine” appeared as the most important function. In addition to being fulfilled by a highest number of top-priority species (11.3±2.9), this function displayed the highest total species richness (44.6±1.7) and highest functional score (91.0±6.8) (Fig 4b). In decreasing order, “Food”, “Sale”, “Energy”, “Fodder” and “Craft” followed “Medicine” in terms of importance.

In the PCA-biplot, the first two axes accounted for 67.6% of observed variation (Table 2) and thus were used to describe the relationships between tree functions and woody species. The first axis was positively and significantly correlated with “Sale”, “Food” and “Energy”, while the second axis was positively and significantly correlated with “Fodder” and negatively correlated with “Medicine” (Table 2).

**Table 2.** Description of the PCA dimensions by their correlation coefficients with the tree functions listed by the discussion group participants in the four villages (Cassou, Dao, Kou and Vrassan) in Ziro province of Burkina Faso

| Variable<br>(Tree function) | Axis 1 (38.7%) |  | Axis 2 (28.9%) |  |
| --- | --- | --- | --- | --- |
|  | Correlation | *p* | Correlation | *p* |
| Energy | 0.56 | 0.021 | -- | -- |
| Fodder | -- | -- | 0.84 | <0.001 |
| Food | 0.75 | <0.001 | 0.43 | 0.022 |
| Medicine | 0.46 | 0.014 | -0.65 | <0.001 |
| Sale | 0.92 | 0.041 | -- | --- |

Furthermore, it revealed relationships between “Food” and “Sale”, and between “Energy” and “Medicine” with “Fodder” representing the third isolated group (Fig 5). Axis 1 was driven by the association of “Sale”, “Food”, “Energy”, and “Medicine” with species such as *V. paradoxa, P. biglobosa, Detarium microcarpum, Acacia macrostachya, Adansonia digitata* and *T. indica*. On the other hand, *B. costatum, A. africana, Balanites aegyptiaca* and *Strychnos spinosa* tended to cluster with “Fodder”, the primary contributing factor in the construction of Axis 2.

**Fig 5.**
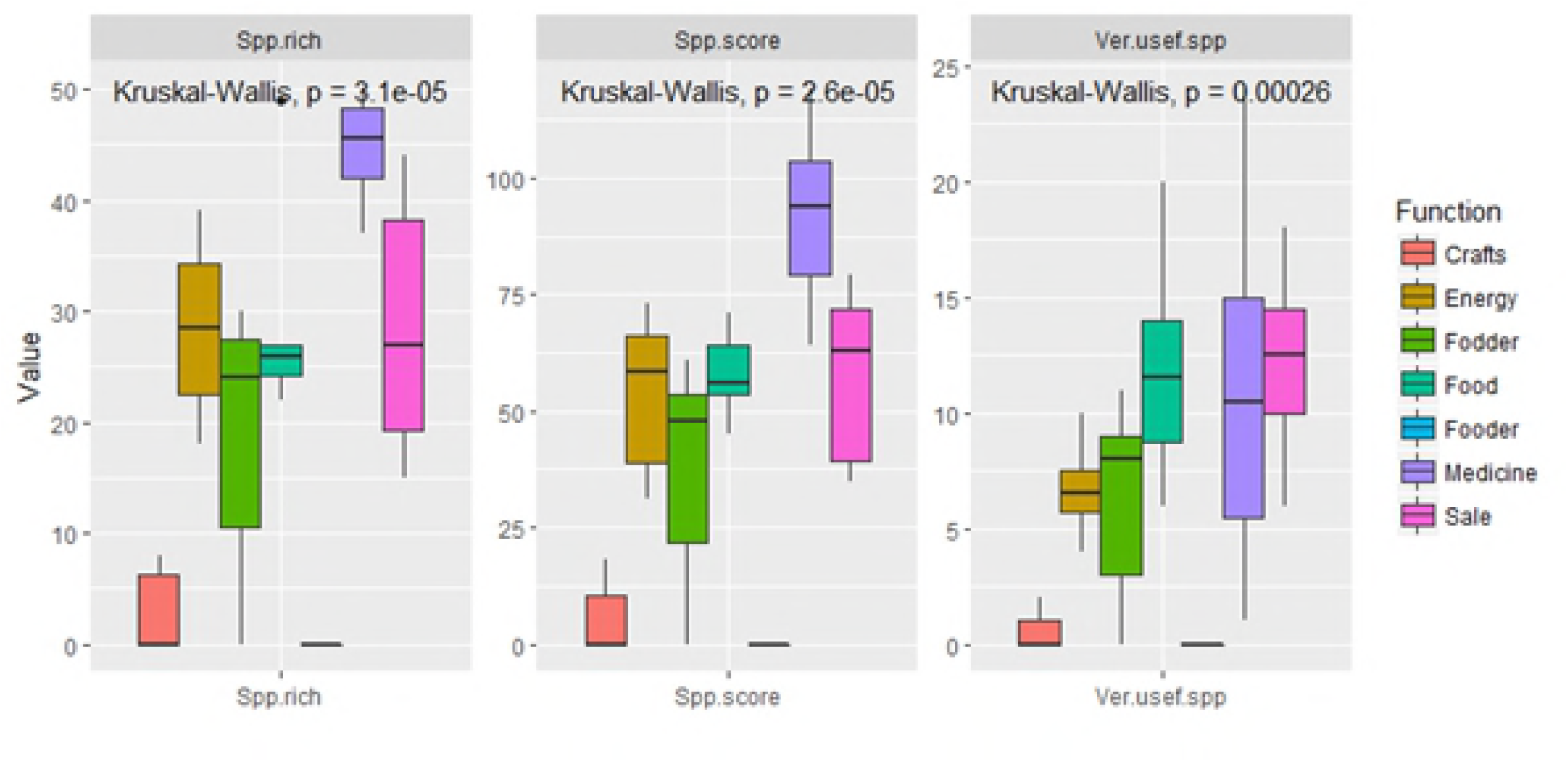
Biplot of the Principal Component Analysis assessing the relationship between woody species and attributes assigned by the four villages (Cassou, Dao, Kou and Vrassan) in Ziro province of Burkina Faso. Afzeafri: *Afzelia africana*, Balaaegy: *Balanites aegyptiaca*, Stryspin: *Strychnos spinosa*, Faidalbi: *Faidherbia albida*, Kayasene: *Khaya senegalensis*, Ficugnap: *Ficus gnaphalocarpa*, Sclebirr: *Sclerocarya birrea*, Termavic: *Terminalia avicennioides*, Cappcory: *Capparis corymbosa*, Acadudg: *Acacia dudgeonii*, Diosmesp: *Diospyros mespiliformis*, Seculong: *Securidaca longipedunculata*, Ficucape: *Ficus capensis*, Casssieb: *Cassia sieberiana*, Burkafri: *Burkea africana*, Ximeamer: *Ximenia americana*, Anogleio: *Anogeisus leiocarpa*, Crosfebr: *Crossopteryx febrifuga*, Lannmicr: *Lannea microcarpa*, Vitedoni: *Vitex doniana*, Ptererin: *Pterocarpus erinaceus*, Acamacr: *Acacia macrostachya*, Adandigi: *Adansonia digitata*, Tamaindi: *Tamarindus indica*, Detamicr: *Detarium microcarpum*, Parkbigl: *Parkia biglobosa*, Vitepara: *Vitellaria paradoxa*.

The top species, with the highest ranking in each village and for each gendered group, are presented in Table 3. The top 28 species listed by respondents were present in all villages, but listing the species and species preferences varied according to gender and from one village to another. Men and women together in Vrassan named the most species, with 20 out of 28 listed as preferred.

**Table 3.**
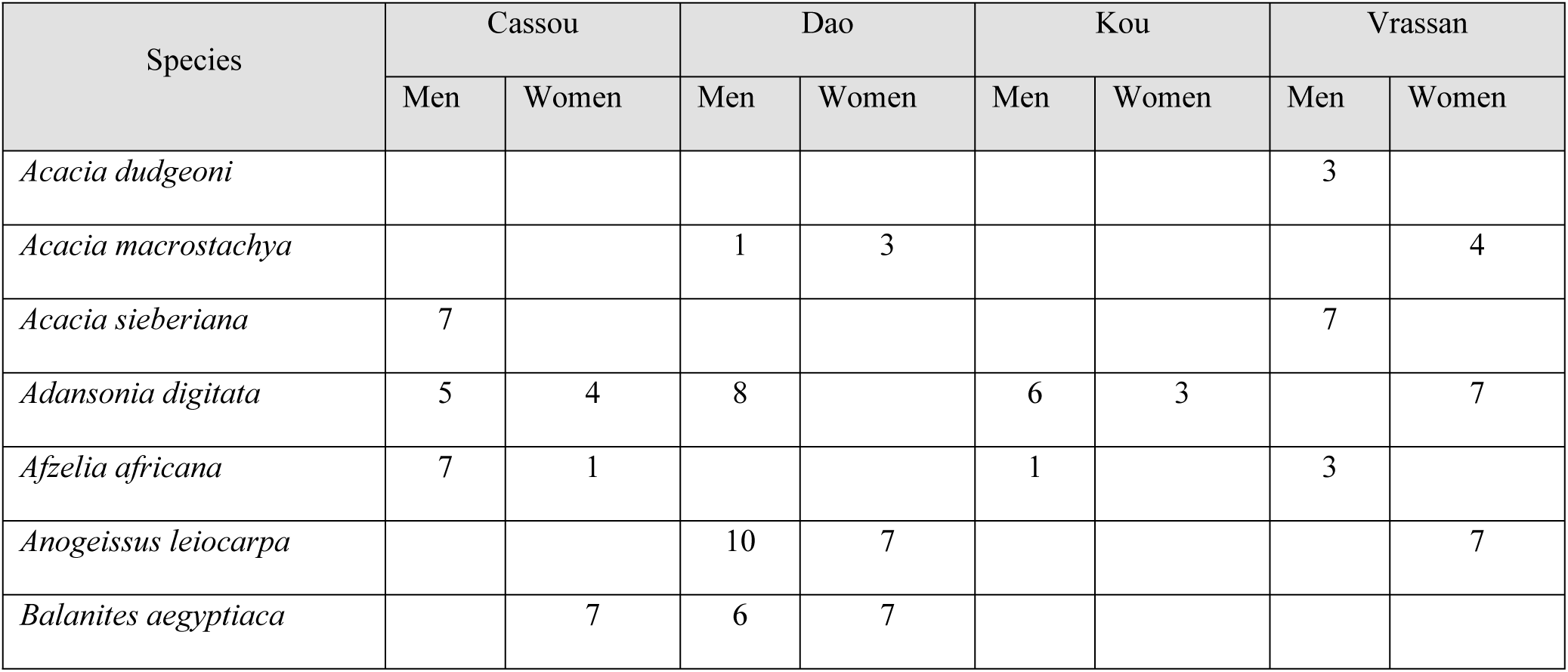
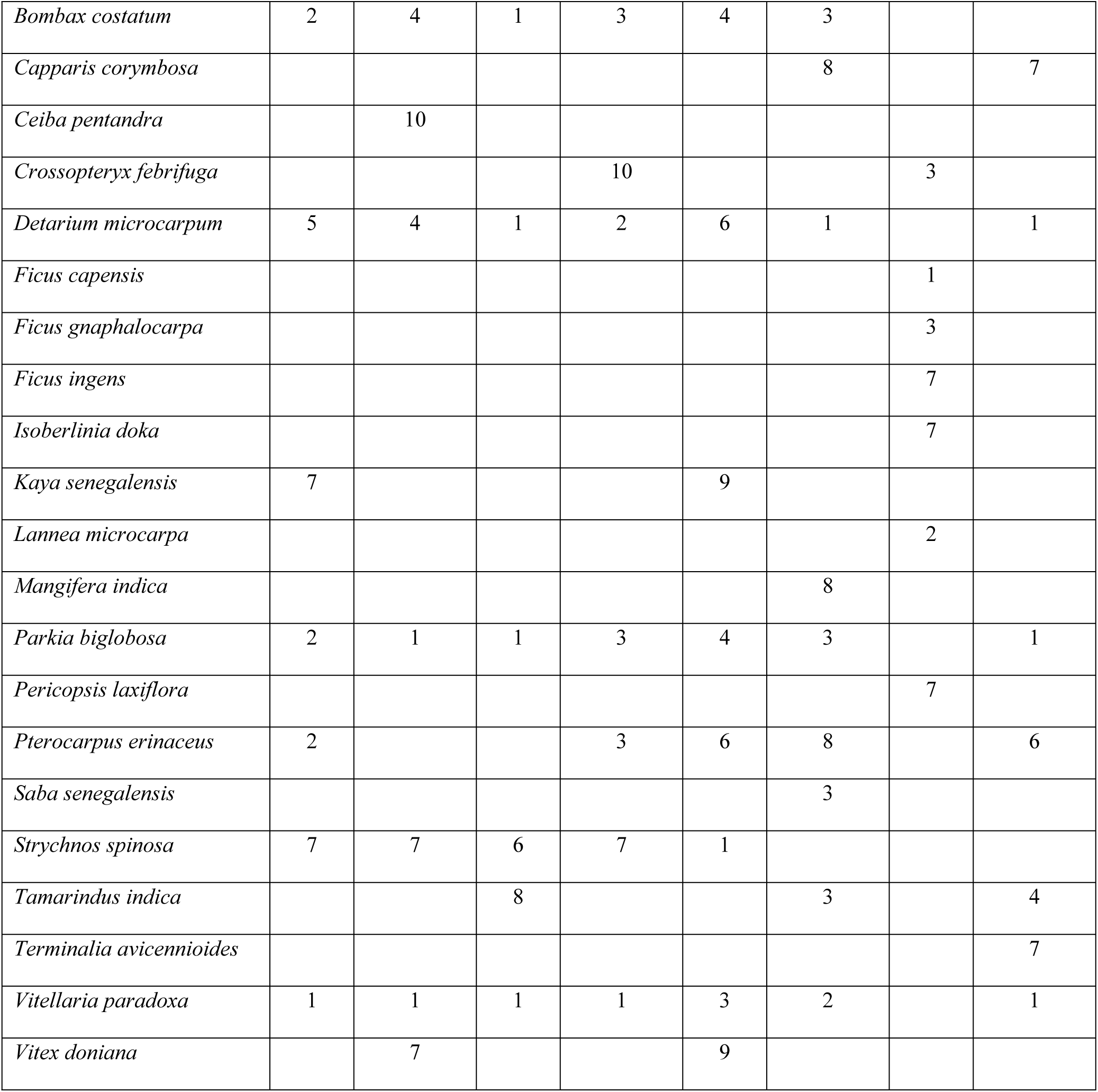
Scoring of the preferred tree species according to the gender in each village (Cassou, Dao, Kou and Vrassan) in Ziro province of Burkina Faso

*V. paradoxa* was the favorite species across the four villages, closely followed by *P. biglobosa*, *B. costatum*, *D. microcarpum, A. digitata* and *S. spinosa*. These species were scored in the top ten in three villages. Four species (*T. indica, B. costatum*, *A. africana* and *S. spinosa*) were cited in three villages, and eight species (*Capparis corymbosa, Vitex doniana, Khaya senegalensis, Crossopteryx febrifuga, Balanites aegyptiaca, Anogeissus leiocarpa, Acacia sieberiana* and *Acacia macrostachya*) were commonly preferred in two villages. Eleven species (*Terminalia avicennioides, Saba senegalensis, Pericopsis laxiflora, Mangifera indica, Lannea microcarpa, Isoberlinia doka, Ficus ingens, Ficus gnaphalocarpa, Ficus capensis, Ceiba pentandra* and *Acacia dudgeonii*) were listed only in one village. It is noteworthy that only one of the 28 species (*Mangifera indica*) is exotic to Burkina Faso. The general interest in the 28 species revealed by the discussion groups augurs well for local tree species domestication initiatives.

### Gender differentiated species preferences

Although some noticeable differences appeared among the measured variables, the use of univariate statistics revealed no significant difference for any variable between the study villages probably due to high coefficients of variation ranging from 49.1 to 136.4 % (Table 4).

**Table 4.** Mean values and coefficient of variation of very useful tree species, species richness and tree function score according to men and women across the villages (Cassou, Dao, Kou and Vrassan) in Ziro province of Burkina Faso

|  | Very useful species |  |  |  | Species richness |  |  |  | Tree function score |  |  |  |
| --- | --- | --- | --- | --- | --- | --- | --- | --- | --- | --- | --- | --- |
|  | Men | CV (%) | Women | CV (%) | Men | CV (%) | Women | CV (%) | Men | CV (%) | Women | CV (%) |
| Global | 8.1±1.1a | 68.9 | 7.8±1.4a | 86.0 | 27.3±3.2a | 58.4 | 22.2±3.1a | 68.2 | 53.8±6.6a | 59.8 | 45.5±6.2a | 67.1 |
| Cassou | 6±1.5a | 62.4 | 8.5±2.1a | 60.0 | 30.5±7.2a | 57.8 | 29.8±6.7a | 55.0 | 57.2±13.4a | 57.6 | 52.5±10.5a | 49.1 |
| Dao | 8.5±2.3a | 67.3 | 10.8±3.5a | 78.4 | 23±6.6a | 70.2 | 22.7±6.5a | 70.4 | 46.8±14.1a | 73.7 | 54±15.5a | 70.4 |
| Kou | 7.3±1.9a | 62.0 | 5.5±3.1a | 136.4 | 25.3±6.6a | 63.7 | 18.5±5.6a | 74.2 | 50.5±12.3a | 59.4 | 37.3±11.5a | 75.3 |
| Vrassan | 10.7±3.2a | 74.2 | 6.3±2.3a | 88.6 | 30.2±6.9a | 55.7 | 17.6±6.2a | 85.7 | 60.8±15.6a | 62.7 | 38.2±13.6a | 87.1 |

In contrast, Figure 6 showed that the first two axes of the PCA biplot accounted for a total variance of 67.6%. Woody vegetation attributes such as “Energy” and “Sale” were significantly correlated to Axis 1 of the biplot, indicating that women are strongly related to the collection of wood for fuelwood used for their own needs and for sale of the surplus to generate income (Table 5; Fig 6). The second axis correlated positively and significantly with “Fodder,” showing men’s preferences, attributable to their responsibilities for feeding household livestock.

**Table 5.**
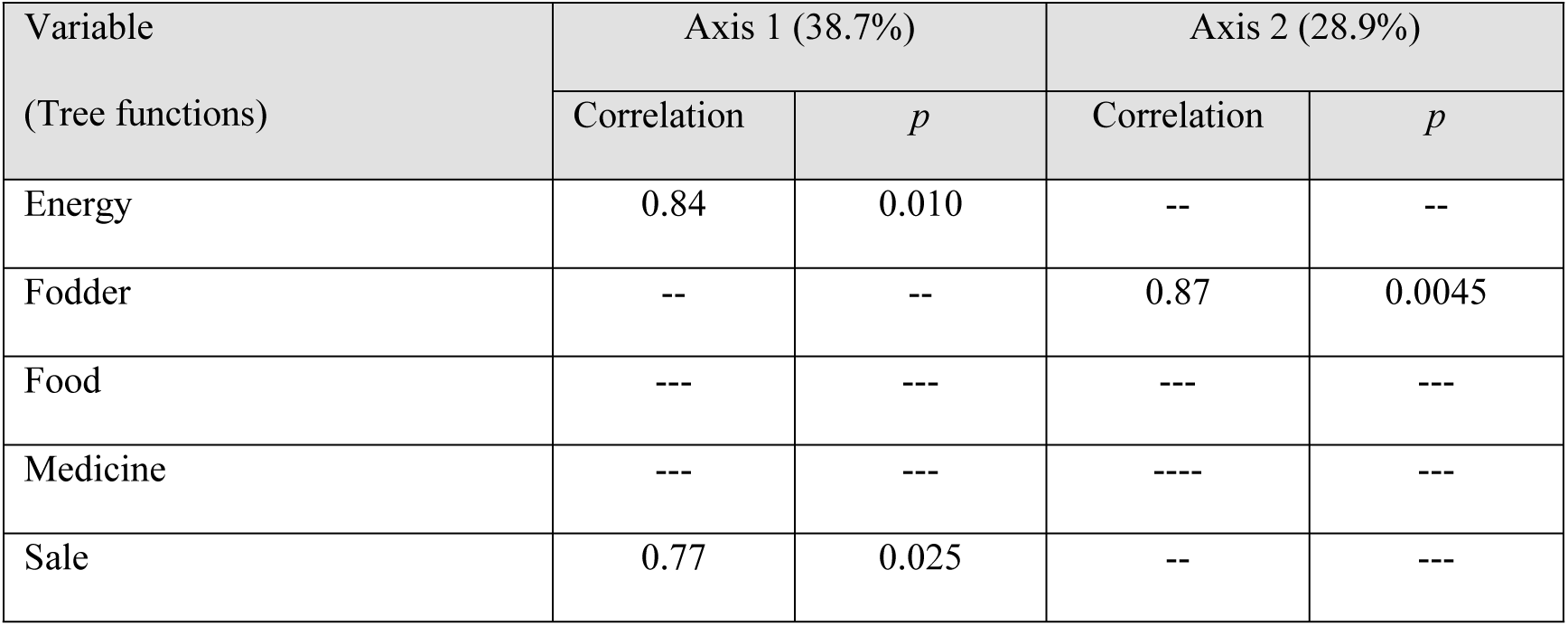
Description of the PCA dimensions by their correlation coefficients with the tree functions listed by the respondents of the four villages (Cassou, Dao, Kou and Vrassan) in Ziro province of Burkina Faso and the value of measured variables (abundance of very useful tree species, tree species richness and total score of tree function).

| Variable<br>(Tree functions) | Axis 1 (38.7%) |  | Axis 2 (28.9%) |  |
| --- | --- | --- | --- | --- |
|  | Correlation | *p* | Correlation | *p* |
| Energy | 0.84 | 0.010 | -- | -- |
| Fodder | -- | -- | 0.87 | 0.0045 |
| Food | --- | --- | --- | --- |
| Medicine | --- | --- | ---- | --- |
| Sale | 0.77 | 0.025 | -- | --- |

**Fig 6.**
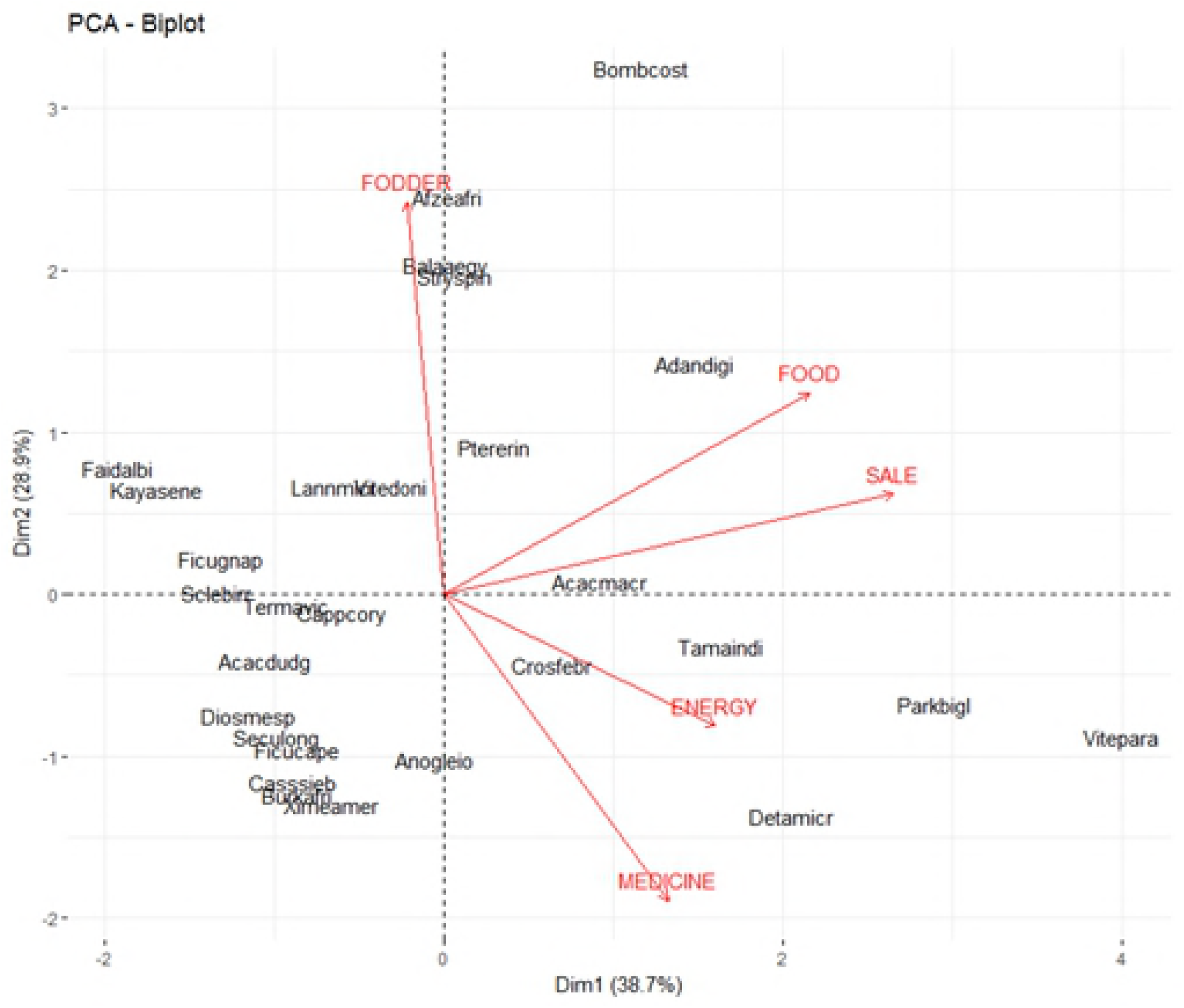
Biplot of the Principal Component Analysis assessing the relationship between gendered groups and woody vegetation attributes in the studied landscape of the four villages (Cassou, Dao, Kou and Vrassan) in Ziro province of Burkina Faso.

### Carbon stock potential for priority woody species

The dbh and height of 1,067 and 1,405 individuals of the preferred woody species were measured in Cassou and Kou, respectively (S2 Table). In the four land use and land cover categories (LULCc) considered in this study, *V. paradoxa,* often associated with *D. microcarpum*, is the species most encountered (Table 6).

**Table 6.** Top five dominant woody species per land use and land cover category in Cassou and Kou (values in parentheses correspond to the total number of individuals)

| Site | Cropland | Shrubland | Woodland | Forest |
| --- | --- | --- | --- | --- |
| Cassou | *Vitellaria paradoxa* (52) | *Vitellaria paradoxa* (71) | *Vitellaria paradoxa* (52) | ---- |
|  | *Parkia biglobosa* (09) | *Detarium microcarpum* (64) | *Detarium microcarpum* (44) | ---- |
|  | *Lannea acida* (04) | *Combretum molle* (37) | *Crossopteryx febrifuga* (35) | ---- |
|  | *Detarium microcarpum* (02) | *Lannea acida* (31) | *Piliostigma thonningii* (31) | ---- |
|  | *Lannea microcarpa* (02) | *Crossopteryx febrifuga* (30) | *Terminalia laxiflora* (27) | ---- |
| Kou | *Vitellaria paradoxa* (272) | *Vitellaria paradoxa* (46) | *Vitellaria paradoxa* (67) | *Anogeisus leiocarpa* (114) |
|  | *Bombax costatum* (22) | *Piliostigma thonningii* (14) | *Detarium microcarpum* (60) | *Detarium microcarpum* (65) |
|  | *Sterculia setigera* (17) | *Detarium microcarpum* (11) | *Anogeisus leiocarpa* (24) | *Vitellaria paradoxa* (42) |
|  | *Terminalia avicenioides* (15) | *Acacia macrostachya* (10) | *Combretum fragrans* (18) | *Acacia macrostachya* (27) |
|  | *Crossopteryx febrifuga* (12) | *Terminalia avicenioides* (10) | *Burkea africana* (15) | *Combretum fragrans* (26) |

The average aboveground carbon stocks (AGC) computed varied from 14.1 ±2.2 to 78.1 ± 5.8 MgC ha^−1^ (±indicates standard error throughout) across the LULCc of Cassou and Kou. The lowest value of AGC was in woodland in Kou (14.1±3.4 MgC ha^−1^) and the highest value in cropland in Kou (78.1 ± 5.8 MgC ha-1). Using a pairwise comparison (Wilcoxon test), we found that there was a significant difference among the amount of AGC stored in shrubland (*p* <0.0001) and woodland of the two sites (*p* <0.0001). However, there was no significant difference in the amount of AGC stored in croplands (p=0.34). Forests were encountered only in Kou and they stored 14.1 ±2.2 MgC ha^−1^ of AGC. As the cultivated land is estimated to be 20.4% and 68.9% in Cassou and Kou, respectively, their respective parkland agroforests could potentially store 1,452.16 Mg C and 5,585.09 Mg C in aboveground biomass.

Among the preferred woody species studied, *V. paradoxa* stored the highest quantity of carbon in the two sampling sites (1,460.6 ±270.9 to 2,798.1±521.0 kg C ha^−1^) while the lowest stocks were found in Cassou with *Strychnos innocua* (8.5±2.5 kg C ha^−1^) and Kou with *Grewia bicolor* (1.6±1.3 kg C ha^−1^). In Cassou, the highest carbon stocks were stored by *V. paradoxa, A. digitata, P. biglobosa, D. microcarpum*, *C. febrifuga, Piliostigma thonningii, Ficus sycomorus, Isoberlinia doka and Terminalia laxiflora* while *V. paradoxa, T. indica, A. leiocarpa, B. costatum, P. erinaceus, A. africana, C. febrifuga* and *D. microcarpum* displayed the greatest values in Kou (S3 Table). *V. paradoxa* stored the largest percentage of carbon in Cassou (32.9 %) and Kou (55.3%), followed by *D. microcarpum* (6.4%) and *P. biglobosa* (4.7%) in Cassou, *A. leiocarpa* (5.4%) and *C. febrifuga* (3.7%) in Kou (Fig 7).

**Fig 7.**
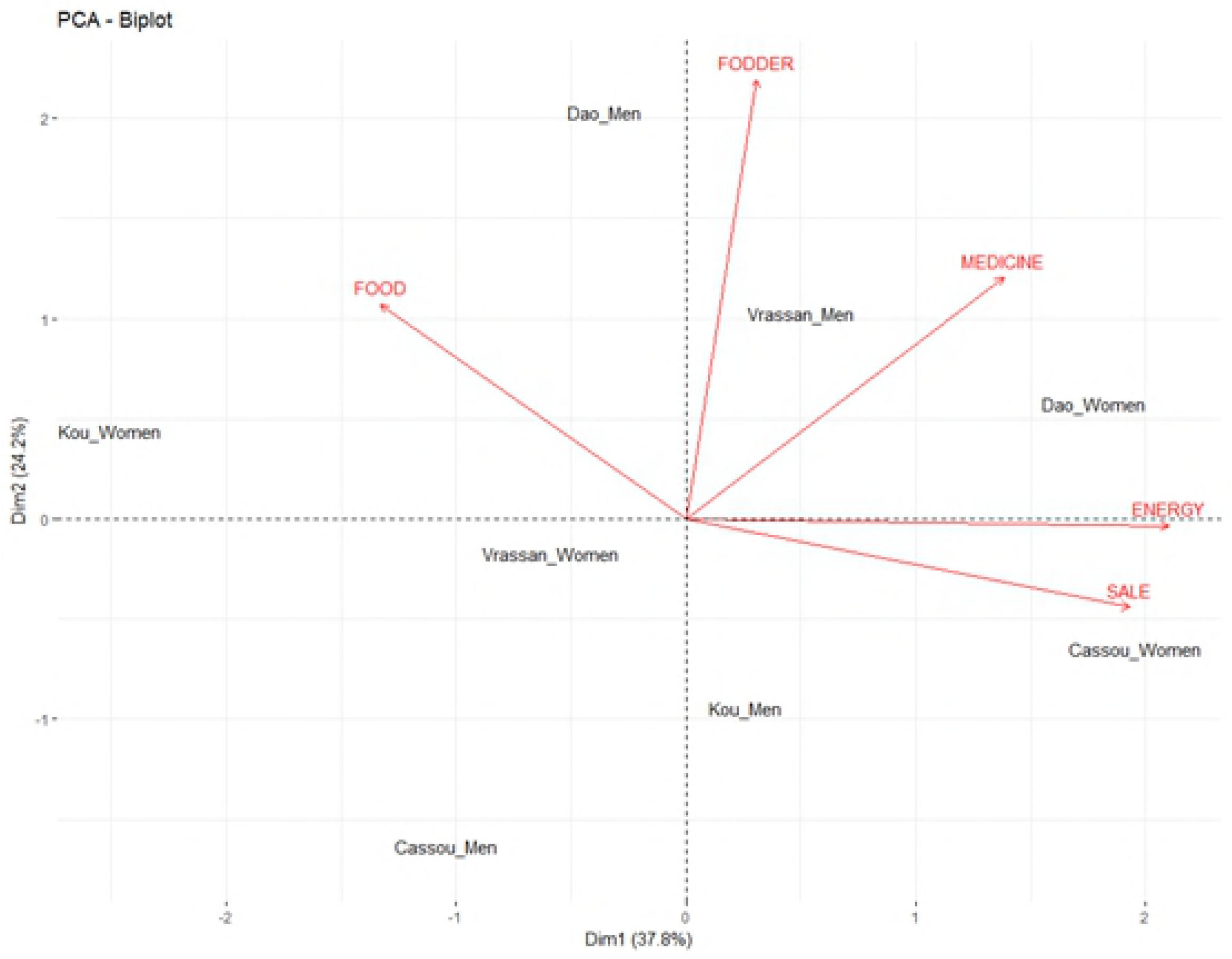
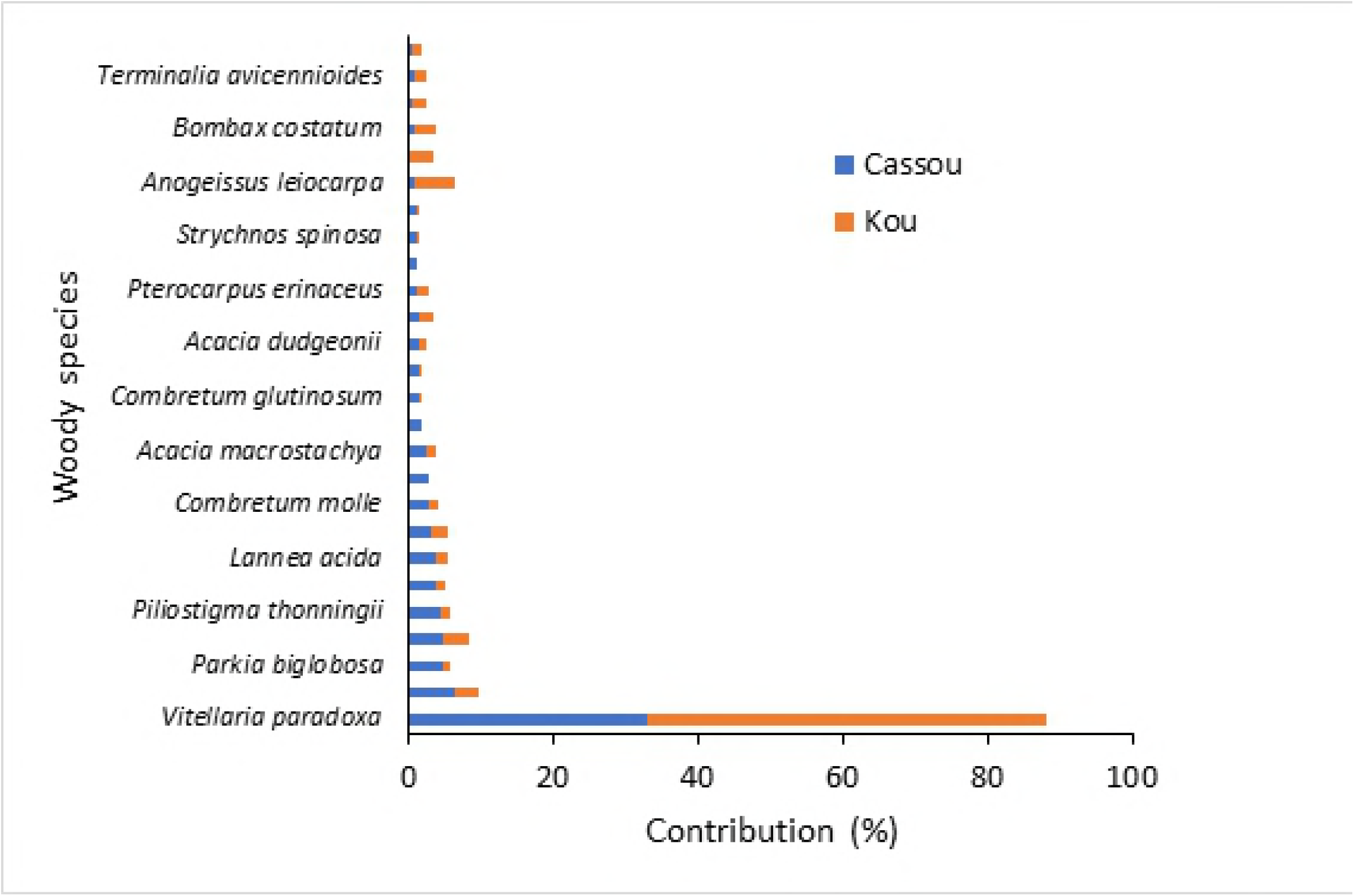
Contribution of preferred woody species to the pool of carbon stock in Cassou and Kou in Ziro Province of Burkina Faso.

The lowest carbon stocks were recorded for *Lannea microcarpa* and *B. aegyptiaca* in Kou, and for *T. indica* and *Khaya senegalensis* in Cassou. The Wilcoxon test statistics showed significant differences in the carbon stored by the preferred woody species in Cassou and Kou (*p* <0.0001). The potential of AGC stored by the preferred woody community was estimated at 5,766.17 Mg C (95% CI: [5,258.2; 6,274.2 Mg C]) in Kou and 6,664.0 Mg C (95% CI: [5,810.2; 7,517.8 Mg C]) in Cassou.

## Discussion

### Functions of smallholders preferred woody species

The overall species richness listed by respondents in this study accounts for 29.3% of all native woody species in Burkina Faso, reported in a previous study as 376 species (including 96 exotic species), 214 genera and 55 families [30]. More than 98% of those listed by the respondents are indigenous species that farmers protect or nurture in their farms. The high species richness number underscores the importance of these species to the rural communities who rely on them to sustain their livelihoods. This finding confirms information from other studies about the critical role of traditional agroforestry practices in supporting biodiversity through *in situ* conservation of tree species [47-48]. This number of woody species cited by respondents also is indicative of the extent to which local people serve as a repository of knowledge on their local vegetation. Previous studies reported that local knowledge can provide important information on plant species richness, diversity, abundance and rarity, necessary for understanding not only local vegetation dynamics but also fundamental management strategies for sustainable use and conservation of natural vegetation [49-50].

Among the top 28 woody species that respondents listed in this study, 18 were identified as top-priority because of their role in providing products and services to sustain livelihoods (“Food”, “Energy”, “Sale”, “Fodder”) and health status (“Medicine”) of smallholders. Local communities prefer most species identified as priority owing to their multipurpose functions providing several products and services to local people, including many income sources. This confirms the statement of Faye et al. [51] about the yearly contribution of agroforestry products, as high as 650 USD, in sustaining household livelihoods in the Sahel. For instance, local communities highly appreciate the foods and condiments provided by *V. paradoxa, P. biglobosa, T. indica* and *A. digitata*. Previous studies have reached similar conclusions about the socioeconomic importance of these species in West Africa [52-55]. Women greatly value *V. paradoxa* nuts for their high oil content, and in all four villages they listed *A. digitata* as top-priority because the leaves are an important sauce ingredient consumed throughout the year, also reported in Faye et al. [28] for smallholders in Mali, Burkina Faso, Niger and Senegal. The critical role *A. africana, P. erinaceus,* and *B. aegyptiaca* play in providing fodder elicited men’s top-priority ranking. This finding accords with that of Naah et al. [56] who reported that men are repository of local knowledge on forage. These trees also supply farmers with fuelwood, as well as several other services [57-59].

Registering “Medicine” (96%) and “Food” (94%) as top-priority functions indicates smallholders’ knowledge of trees for nutritional and health properties. Smallholders’ reliance on more accessible and affordable herbal medicines for health care illustrates the major challenges that health care presents in Burkina Faso [60]. These categories were followed by “Sale” (87%), “Energy” (56%), and “Fodder” (45%), consistent with the prime importance of these functions for household income generation, fuel and livestock feed [28, 51]. The study revealed a gender-oriented preference for tree species and functions in all study villages. Overall, while men focused on provisioning services to feed their livestock, women focused on energy (fuel wood) and income.

The positive relationships among most woody species’ functions indicate their multiple uses for smallholders, generally true throughout the West African Sahel [28]. Some fruits, fats, leafy vegetables, nuts and condiments may be used as food, to cure certain diseases [28], and sold for cash. The confirmation that farmers value functional diversity over species richness is important to the success of biocarbon projects which aim at alleviating rural poverty and strengthening the resilience of the parkland agroforests in the Sahelian and Sudanian ecozones of West Africa.

### Carbon stock potential of community preferred woody species

Findings revealed that *V. paradoxa, P. biglobosa, D. microcarpum, A. leiocarpa, C. febrifuga* and *B. costatum* stored more carbon than the other preferred woody species in Kou and Cassou. This could be explained by the fact that these species were abundant and their dbh was larger than the dbh of the other preferred woody species. Apart from *A. leiocarpa* and *B. costatum* which are commonly found in natural vegetation, *V. paradoxa, P. biglobosa,* and *D. microcarpum* are encountered in parklands. *V. paradoxa* and *P. biglobosa* are two of the dominant tree species of parklands in Burkina Faso [62]. They have great socioeconomic importance for rural communities as well as ecological functions, providing a range of ecosystem services, including shade and habitat for biodiversity, and mitigative services by storing carbon [5, 28]. The capacity of these two species to supply goods and services to the local people confers them both systematic protection against anthropogenic interventions, such as excessive harvesting and pruning, and enhanced restoration through farmer-managed natural regeneration (FMNR).

Carbon stocks varied greatly among woody species owing to the different population densities per cluster and their morphological differences. In addition, there were large differences in carbon stocks between the two sites for species like *V. paradoxa*, *A. leiocarpa*, *B. costatum*, *Pericopsis laxiflora*, and *P. biglobosa*, possibly due to factors that were not assessed in this study, such as microclimate, soil, geomorphology or genetic differences. The carbon stock obtained in Kou for *A. leiocarpa*, for example, is far lower than the 907.9 kg C ha^−1^ reported by Eneji et al. [63] in a Forest Reserve of Benue State, Nigeria, and that stored by *P. biglobosa* in the same forest (301.6 kg C ha^−^1) is slightly lower that that measured in Cassou (372.3±111.0 kg C ha^−1^). The estimated total carbon stored in studied parklands in Cassou (1,452.2 Mg C) and Kou (5,585.1 Mg C) is lower than the amount estimated for mature parklands of the West Afican Sahel (1,284 Tg: [19]).

Three reasons could explain the lower values obtained in our study. First, measuring only trees with heights ≥3m reduced the number of inviduals considered. Second, the use of average genus and family wood density in the absence of species wood density could have been a source of error as wood density of species within the same genus can differ greatly, and even more so within the same family [64]. Third, due to higher land-use pressure in the study sites as compared to the sites of previous studies (e.g Forest Reserve of Benue State), most of the trees are thinned. The aggregated impact of these sources of errors could have resulted in much lower carbon stock values for each species and for the overall landscapes. The significantly higher stock of carbon in trees in Kou relative to Cassou is likely due to Kou’s high density of *V. paradoxa* and *A. leiocarpa.* Cassou has much better market access, intensifying anthropogenic pressure on woody species there as opposed to Kou. Furthermore, among the land use categories considered in this study, the highest overall mean AGC stock was recorded for croplands. This can be attributed to the fact that croplands in the two sites were mainly composed of large *V. paradoxa* trees, and *V. parado*xa was the most abundant species amongst all the woody species measured in both sites. The AGC stocks recorded in woodland and shrubland were higher in Cassou compared to Kou. This difference could be explained by the management system of the two sites and the variability of dendrometric parameters in the individuals of the same species for different geographical locations and ages [65].

The practical implications of our findings are of four-folds. The most likely approach is to (i) recommend the preferred tree and shrub species for FMNR on farmed lands; (ii) plant the preferred species in degraded parklands using selected germplasm to increase their production over time; (iii) train smallholders in tree domestication techniques to promote the species with high carbon sequestration potential; and (iv) promote afforestation and reforestation within the degraded portions of Cassou Protected Forest. Because the climate is becoming hotter and drier and with more variability in rainfall, it is recommended to use germplasm (i.e., seeds, rooted cuttings, etc.) originating from drier locations within the same general region [64].

## Conclusion

This study highlighted the importance of woody species in local people’s strategies to sustain their livelihoods, which may be instrumental to climate change adaptation. It was revealed that local people in the four villages have good knowledge of and use woody species (trees and shrubs) for several functions, but knowledge of woody species varies according to species’ attributes and smallholders’ gender. For biocarbon initiatives, it is recommended to use *V. paradoxa*, *A. leiocarpa*, *P. biglobosa*, and *D. microcarpum* owing to their appreciable carbon storage potential. The challenge now is to determine a cost-effective way to involve local men and women together with technical teams from research institutes and universities in participatory tree domestication and smart land use programs which rely on innovative smallholder-based extension approaches.

## Acknowledgments

This study was funded by the Ministry of Foreign Affairs of Finland through the Building Biocarbon and Rural Development in West Africa (BIODEV) project implemented in Burkina Faso, Guinea-Conakry, Mali and Sierra Leone. The authors are thankful to Institut de l’Environnement et de Recherches Agricoles (INERA) of Burkina Faso for their logistic support. They also extend their special thanks to the smallholders from the study villages who willingly accepted to be interviewed despite their busy schedules during the cropping season.

## Supporting information

S1 Dataset. Data for Figures 4, 5 and 6

S2 Dataset. Dendrometric parameters used to compute aboveground biomass and carbon in Cassou and Kou

S1 Table. List of plant species recorded in the four villages (Cassou, Dao, Kou and Vrassan) in Ziro province of Burkina Faso

S2 Table. Structural characteristics of the preferred woody species at the study sites of Cassou and Kou in Ziro province of Burkina Faso

S3 Table. Estimated aboveground carbon stock (Average ± SE) at cluster level in Cassou and Kou in Ziro provience od Burkina Faso

**Author Contributions**
Conceptualization: Jérôme E. Tondoh, Kangbéni Dimobe, Data curation: Kangbéni Dimobe, Jérôme E. Tondoh
Formal analysis: Kangbéni Dimobe.
Funding acquisition: Antoine Kalinganire, Jules Bayala, Jérôme E. Tondoh, John C. Weber, Methodology: Jérôme E. Tondoh, John C. Weber, Kangbéni Dimobe,
Software: Kangbéni Dimobe
Writing - original draft: Jérôme E. Tondoh, Kangbéni Dimobe, Writing - review & editing: Kangbéni Dimobe, Jérôme E. Tondoh, John C. Weber, Jules Bayala, Karen Greenough, Antoine Kalinganire

## References

1. Boffa J-MJ. West African agroforestry parklands: keys to conservation and sustainable management. Unasylva; 2000. pp. 11–17.

2. Lamien N, Boussim JI, Nygard R, Ouédraogo JS, Odén PC, Guinko S. Mistletoe impact on Shea tree (*Vitellaria paradoxa* CF Gaertn.) flowering and fruiting behaviour in savanna area from Burkina Faso. Environ Exp Bot. 2006; 55: 142–148.

3. Ouedraogo B. Household energy preferences for cooking in urban Ouagadougou, Burkina Faso. Energy Policy. 2006; 34: 3787–3795.

4. Coulibaly-Lingani P, Tigabu M, Savadogo P, Oden PC, Ouadba JM. Determinants of access to forest products in southern Burkina Faso. For Policy Econ. 2009; 11: 516–524.

5. Bayala J, Sanou J, Teklehaimanot Z, Kalinganire A, Ouédraogo JS. Parklands for buffreing climate risk and sustaining agricultural production in the Sahel of West Africa. Curr Opin Environ Sustain. 2014; 6: 28–34.

6. Sinare H, Gordon JL. Ecosystem services from woody vegetation on agricultural lands in Sudano-Sahelian West Africa. Agric Ecosyst Environ. 2015; 200: 186–199.

7. Sinare H, Gordon JL, Kautsky EE. Assessment of ecossyem services and benefits in village landscapes - A case study from Burkina Faso. Ecosyst Serv. 2016; 21: 141–152.

8. Dimobe K, Goetze D, Ouédraogo A, Mensah S, Akpagana K, et al. Aboveground biomass allometric equations and carbon content of the shea butter tree (*Vitellaria paradoxa* CF Gaertn., Sapotaceae) components in Sudanian savannas (West Africa). Agroforestry systems. 2018; 1–14. doi 10.1007/s10457-018-0213-y

9. Ouédraogo I, Nacoulma B M I, Hahn K, Thiombiano A. Assessing ecosystem services based on indigenous knowledge in south-eastern Burkina Faso (West Africa). International Journal of Biodiversity Science, Ecosystem Services & Management. 2014; 10(4): 313–321.

10. Maranz A. Tree mortality in the African Sahel indicates an anthropogenic ecosystem displaced by climate change. J Biogeogr. 2009; 36: 1181–1193.

11. Dimobe K, Ouédraogo A, Soungalo S, Goetze D, Porembski S, Thiombiano A Identification of driving factors of land degradation and deforestation in the Wildlife Reserve of Bontioli (Burkina Faso, West Africa). Glob Ecol Conserv. 2015; 4: 559–571.

12. Etongo D, Djenontin INS, Kanninen M, Fobissie K. Smallholders’ tree planting activity in the ziro province, southern Burkina faso: Impacts on livelihood and policy implications. Forests. 2015; 6: 2655–2677.

13. Berry N, Cross H, Ridell M, Mbow C, Ouédraogo I, Tondoh J. Community biocarbon projects in West Africa: challenges and lessons learned. Report by Bioclimate and the World Agroforestry Centre. 2016; Draft 0.1.

14. Scherr JS Shames S, Friedman R. From climate-smart agriculture to climate-smart landscapes. Agriculture & Food Security. 2012; 1: 1–15.

15. World Bank. The Biocarbon Fund Initiative for sustainable forest landscape. Annual Report; 2016a.

16. World Bank. Biocarbon Fund. Carbon Finance at the World Bank; 2016b.

17. Mahanty S, Suich H, Tacconi L. Access and benefits in payments for environmental services and implications for REDD+: lessons from seven PES schemes. Land use policy. 2013; 31: 38–47.

18. Foster K, Neufeldt H. Biocarbon projects in agroforestry: Lessons from the past for future development. Curr Opin Environ Sustain. 2014; 6: 148–154.

19. Luedeling E, Neufeldt H. Carbon sequestration potential of parkland agroforestry in the Sahel. Clim Change. 2012; 115: 443–461.

20. Reij C, Tapan G, Smale M. Agroenvironmental transformation in the Sahel. Another kind of “Green Revolution”. 2020 Vision Initiative; 2009.

21. Weber CJ, Sotelo Montes C, Abasse T, Sanquetta RC, Silva AD, et al. Variation in growth, wood density and carbon concentration in five tree and shrub species in Niger. New Forest. 2018; 49: 35–51.

22. ICRAF. Building Biocarbon And Rural Development in West Africa (BIODEV). Word Agroforestry Centre. Nairobi, Kenya; 2014.

23. Coulibaly-Lingani P, Savadogo P, Tigabu M, Oden PC. Factors influencing people’s participation in the forest management program in Burkina Faso, West Africa. For Policy Econ. 2011; 13: 292–302.

24. Fontès J, Guinko S. Vegetation map and land use in Burkina Faso. Explanatory note: French Ministry of Cooperation: Campus Project (88313101); 1995.

25. Driessen P, Deckers J, Spaargaren O, Nachtergaele F. Lecture notes on the major soils of the world. Food and Agriculture Organization (FAO); 2001.

26. INSD. Résultats préliminaires du recensement général de la population et de l’habitat de 2006. Ouagadougou, Burkina Faso; 2007.

27. Ouedraogo I, Mbow C, Balinga M, Neufeldt H. Transitions in Land Use Architecture under Multiple Human Driving Forces in a Semi-Arid Zone. Land. 2015; 4: 560–577.

28. Faye MD, Weber JC, Abasse TA, Boureima M, Larwanou M, et al. Farmers’ Preferences for Tree Functions and Species in the West African Sahel. Forests, Trees and Livelihoods. 2011; 20: 113–136.

29. Dagnelie P. Statistique théorique et appliquée. Tome1. Bruxelles, Belgique, De Boeck et Larcier; 1998. pp. 1–517.

30. Thiombiano A, Schmidt M, Dressler S, Ouédraogo A, Hahn K, Zizka G. Catalogue des plantes vasculaires du Burkina Faso. Boissiera. 2012; 65:1–391, ISSN: 0373-2975

31. Zizka A, Thiombiano A, Dressler S, Nacoulma BM, Ouedraogo A, et al. Traditional plant use in Burkina Faso (West Africa): a national-scale analysis with focus on traditional medicine. J Ethnobiol Ethnomed. 2015; 11 (1): 9.

32. Nolan JM, Robbins MC. Cultural conservation of medicinal plant use in the Ozarks. Hum Organ; 1999. pp. 67–72.

33. Vagen T, Winoweicki L, Tamene L, Tondoh JE. The Land Degradation Surveillance Framework: Field Guide; ICRAF/CIAT: Nairobi, Kenya; 2013.

34. Sakkatat P. T-saquare sampling. Bang Phra Center J. 1997; 4: 27–30.

35. Sakkatat P. Estimation of number and density, and random distribution testing of important plant species in Ban Pong Forest, Sansai District, Chiang Mai Province, Thailand using T-Square sampling. Maejo International Journal of Science and Technology. 2007; 1: 64–72.

36. Mitchell K. Quantitative Analysis by the Point-Centered Quarter Method; Hobart and William Smith Colleges. Geneva, Switzerland; 2007.

37. FAO. The Miombo in transition: Woodlands and Welfare in Africa. In Environment and Natural Resources Series 8. Rome, Italy; 2005.

38. Tamene L, Mponela P, Sileshi G W, Chen J, Tondoh EJ. Spatial variation in tree density and estimated aboveground carbon stocks in Southern Africa. Forests. 2016; 7: 1–19.

39. Chave J, Réjou-Méchain M, Búrquez A, Chidumayo E, Colgan MS, et al. Improved allometric models to estimate the aboveground biomass of tropical trees. Glob Chang Biol. 2014; 20 (10): 3177–3190.

40. Chave J, Coomes D, Jansen S, Lewis SL, Swenson NG, Zanne AE. Towards a worldwide wood economics spectrum. Ecol Lett. 2009; 12: 351–366.

41. Zanne AE, Lopez-Gonzalez G, Coomes DA, et al. Global wood density database; 2009. Dryad. Available at: http://datadryad.org/handle/10255/dryad.235. Cited 18 May 2018.

42. Réjou-Méchain M, Tanguy A, Piponiot C, Chave J, Hérault B. Biomass: an r package for estimating above-ground biomass and its uncertainty in tropical forests. Methods Ecol Evol. 2017; 8(9): 1163–1167.

43. Tran DB, Dargusch P, Herbohn J, Moss P. Interventions to better manage the carbon stocks in Australian melaleuca forests. Land use policy. 2013; 35: 417–420.

44. Mensah S, Veldtman R, du Toit B, Glèlè Kakaï R, Seifert T. Aboveground Biomass and Carbon in a South African Mistbelt Forest and the Relationships with Tree Species Diversity and Forest Structures. Forests. 2016; 79: 1–17. doi:10.3390/f7040079

45. R Core Team. R: A language and environment for statistical computing. Vienna, Austria: R Foundation for Statistical Computing; 2016. Retrieved from https://www.R-project.org/

46. Zuur A F, Ieno E N, Elphick CS. A protocol for data exploration to avoid common statistical problems. Methods Ecol Evol. 2010; 1(1): 3–14.

47. McNeely JA, Schroth G. Agroforestry and biodiversity conservation - Traditional practices, present dynamics, and lessons for the future. Biodiversity & Conservation. 2006; 15: 549–554.

48. Assogbadjo AE, Glèlè Kakaï R, Vodouhê FG, Djagoun CAMS, Codjia JTC, Sinsin B. Biodiversity and socioeconomic factors supporting farmers’ choice of wild edible trees in the agroforestry systems of Benin (West Africa). For Policy Econ. 2012; 14: 41–49.

49. Lykke AM. Local perceptions of vegetation change and priorities for conservation of woody-savanna vegetation in Senegal. J Environ Manage. 2000; 59: 107–120.

50. Sop T, Oldeland J, Bognounou F, Schmiedel U, Thiombiano A. Ethnobotanical knowledge and valuation of woody plants species: a comparative analysis of three ethnic groups from the sub-Sahel of Burkina Faso. Environment, Development and Sustainability. 2012; 14: 627–649.

51. Faye MD, Weber CJ, Mounkoro B, Dakouo J-M. Contribution of parkland trees to farmers’ livelihoods: a case study from Mali. Dev Pract. 2010; 20: 428–434.

52. Teklehaimanot Z. Exploiting the potential of indigenous agroforestry trees: *Parkia biglobosa* and *Vitellaria paradoxa* in sub-saharan Africa. Agroforestry Systems. 2004; 61: 207–220.

53. Akpona TJD, Akpona HA, Djossa BA, Savi MK, Daïnou K, et al. Impact of land use practices on traits and production of shea butter tree (*Vitellaria paradoxa* C.F. Gaertn.) in Pendjari Biosphere Reserve in Benin. Agroforestry Systems. 2015; 90(4), 607–615.

54. Kébenzikato AB, Wala K, Atakpama W, Dimobe K, Dourma M, et al. Connaissances ethnobotaniques du baobab (*Adansonia digitata* L.) au Togo. BASE. 2015; 19: 247–261.

55. Padakale E, Atakpama W, Dourma M, Dimobe K, Wala K, et al. Woody Species Diversity and Structure of Parkia biglobosa Jacq. Dong Parklands in the Sudanian Zone of Togo (West Africa). Annu Res Rev Biol. 2015; 103–114.

56. Naah JBS, Guuroh RT. Factors influencing local ecological knowledge of forage resources: Ethnobotanical evidence from West Africa’s savannas. J Environ Manage. 2017; 188: 297–307.

57. Sotello Montes C, da Silva DA, Garcia RA, Bolzón de Muñiz GI, Weber JC. Calorific value of *Prosopis africana* and *Balanites aegyptiaca* wood: Relationships with tree growth, wood density and rainfall gradients in the West African Sahel. Biomass Bioenergy. 2011; 35: 346–353.

58. Paré S, Savadogo P, Tigabu M, Ouadba JM, Oden PC. Consumptive values and local perception of dry forest decline in Burkina Faso, West Africa. Environment, Development and Sustainability. 2010; 12: 277–295.

59. Sinsin B, Matig OE, Assogbadjo AE, Gaoué OG, Sinadouwirou T. Dendrometric characteristics as indicators of pressure of *Afzelia africana* Sm. dynamic changes in trees found in different climatic zones of Benin. Biodivers Conserv. 2004; 13: 1555–1570.

60. Mander M. Marketing of indigenous medicinal plants in South Africa: a case study in KwaZulu-Natal; 1998.

61. Kalinganire A, Weber JC, Uwamariya A, Koné B. Improving rural livelihoods through domestication of indigenous fruit trees in the parklands of the Sahel. in Akinnifesi, F.K., Leakey, R.R.B., Ajayi, O.C., Sileshi, G., Tchoundjeu, Z., Matakala, P., Kwesiga, F.R., (eds). Indigenous Fruit Trees in the Tropics: Domestication, Utilization and Commercialization; 2008. pp. 186–203.

62. Bayala J, Heng LK, van Noordwijk M, Ouedraogo SJ. Hydraulic redistribution study in two native tree species of agroforestry parklands of West African dry savanna. Acta Oecol. 2008; 34: 370–378.

63. Eneji SI, Obinna O, Azua TE. Sequestration and carbon storage potential of tropical Forest Reserve and tree species located within Benue State of Nigeria. Journal of Geoscience and Environment Protection. 2014; 2: 157–166.

64. Weber JC, Sotelo Montes C, Kalinganire A, Abasse T, Larwanou M. Genetic variation and clines in growth and survival of *Prosopis africana* from Burkina Faso and Niger: comparing results and conclusions from a nursery test and a long-term field test in Niger. Euphytica. 2015; 205: 809–821.

65. Fearnside PM. Wood density for estimating forest biomass in Brazilian Amazonia. For Ecol Manage. 1997; 90: 59–87.

